## Supplemental figures for "The budding yeast Fkh1 Forkhead associated (FHA) domain promoted a G1-chromatin state and the activity of chromosomal DNA replication origins"

#### Supplementary information

Timothy Hoggard[1], Erika Chacin[2], Allison J. Hollatz[1,3], Christoph F. Kurat[2], and Catherine A. Fox[1,3,4]

[1]Department of Biomolecular Chemistry, School of Medicine and Public Health, University of Wisconsin, Madison [2]Biomedical Center Munich (BMC), Division of Molecular Biology, Faculty of Medicine, Ludwig-Maximilians-Universität in Munich, Martinsried, Germany [3]Integrated Program in Biochemistry, University of Wisconsin, Madison [4]Corresponding author:

### Figure S1

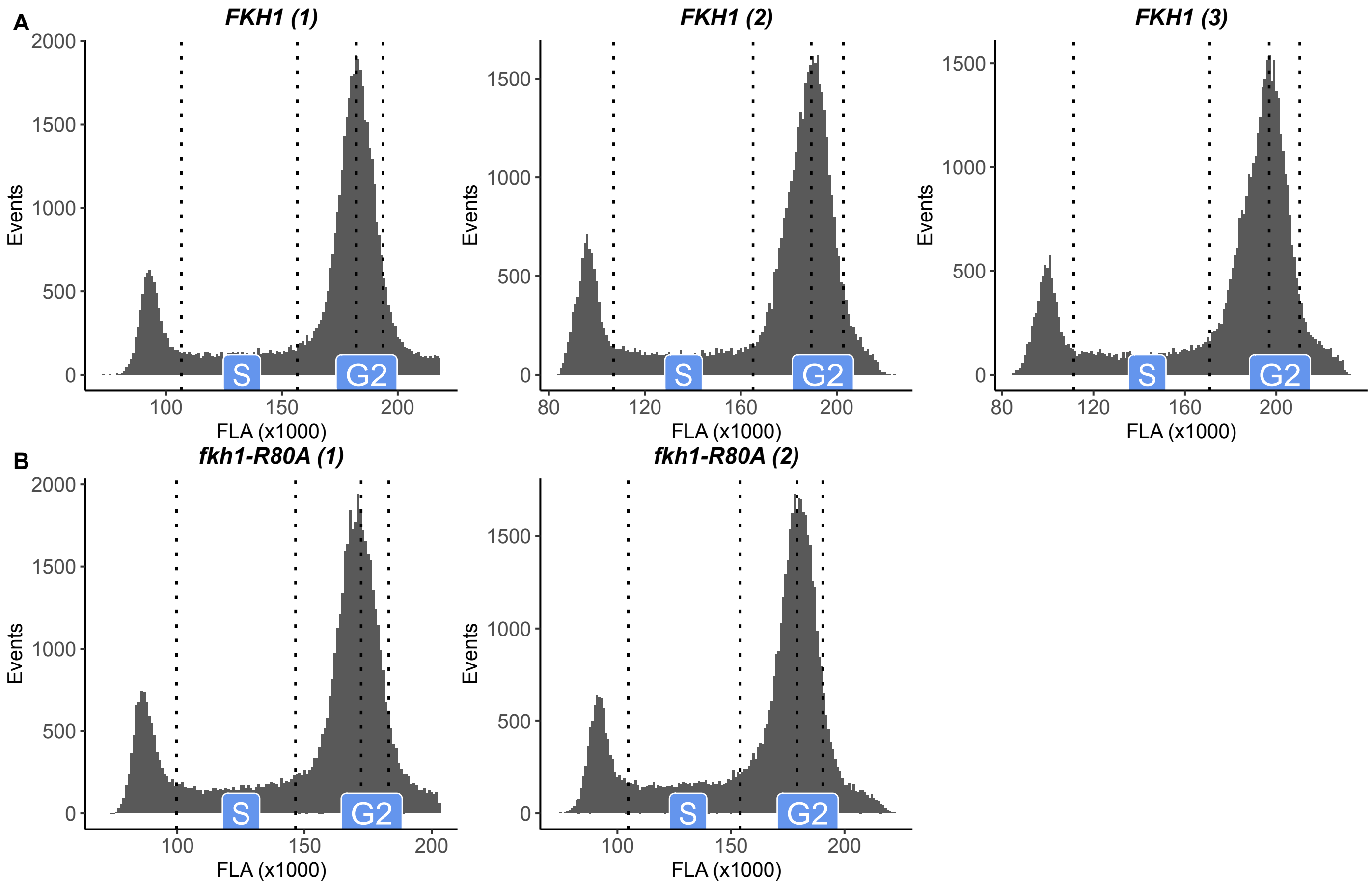


**Figure S1. Defining the S-phase and G2-phase populations used to generate the SortSeq data.** The cell density histograms from the proliferating yeast population as a function of fluorescence are shown for **(A)** Three independent FKH1 yeast and **(B)** Two independent *fkh1-R80A* yeast used to generate the data used in this study. Vertical dotted lines indicate the gates used for the collection of the S-phase and G2-phase populations that were processed for sequencing for SortSeq.

### Figure S2

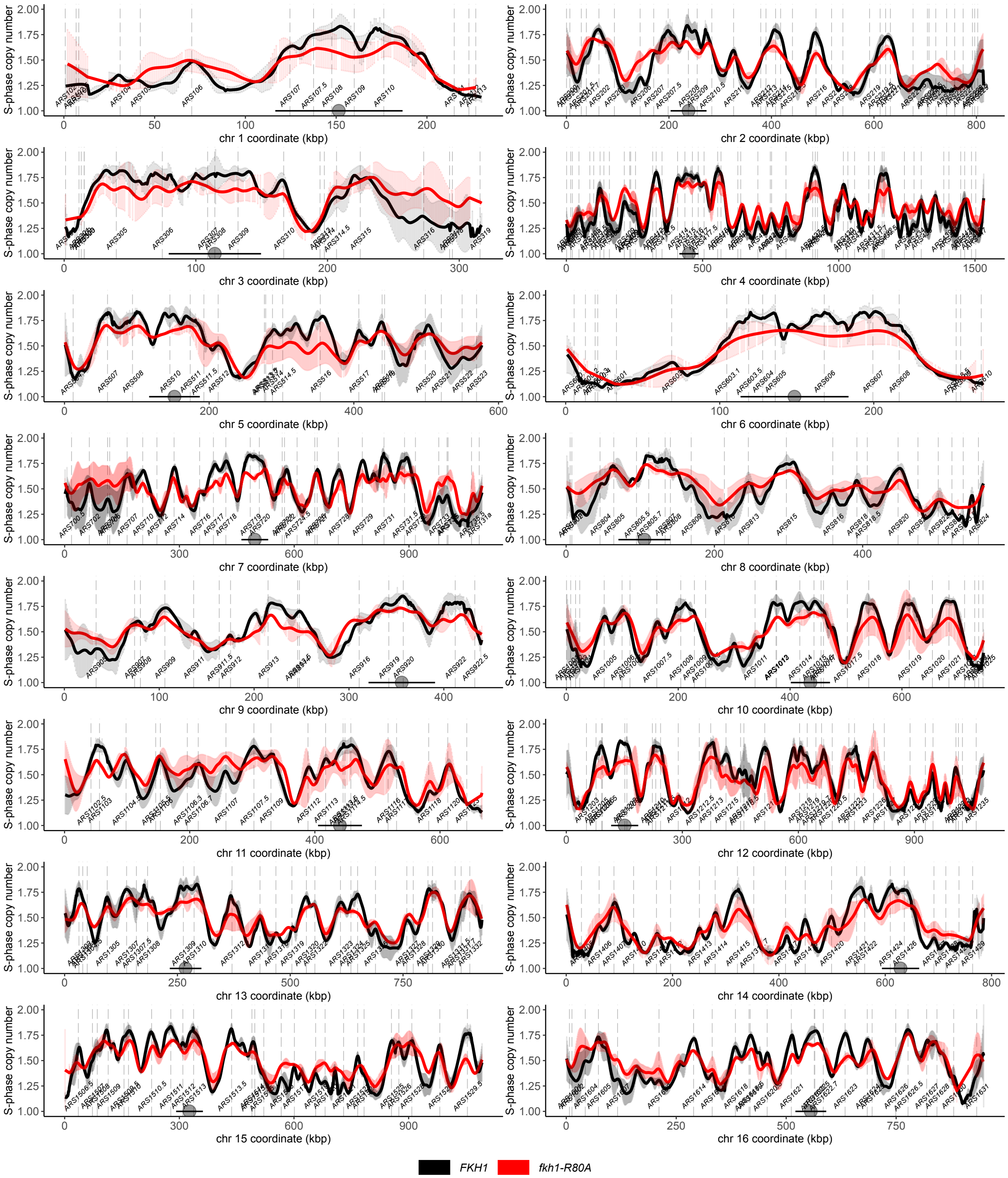


**Figure S2. S-phase SortSeq scans for all yeast nuclear chromosomes.** For details please see the methods, text and Figure legend sections relevant to main text Figure 1A.

### Figure S3

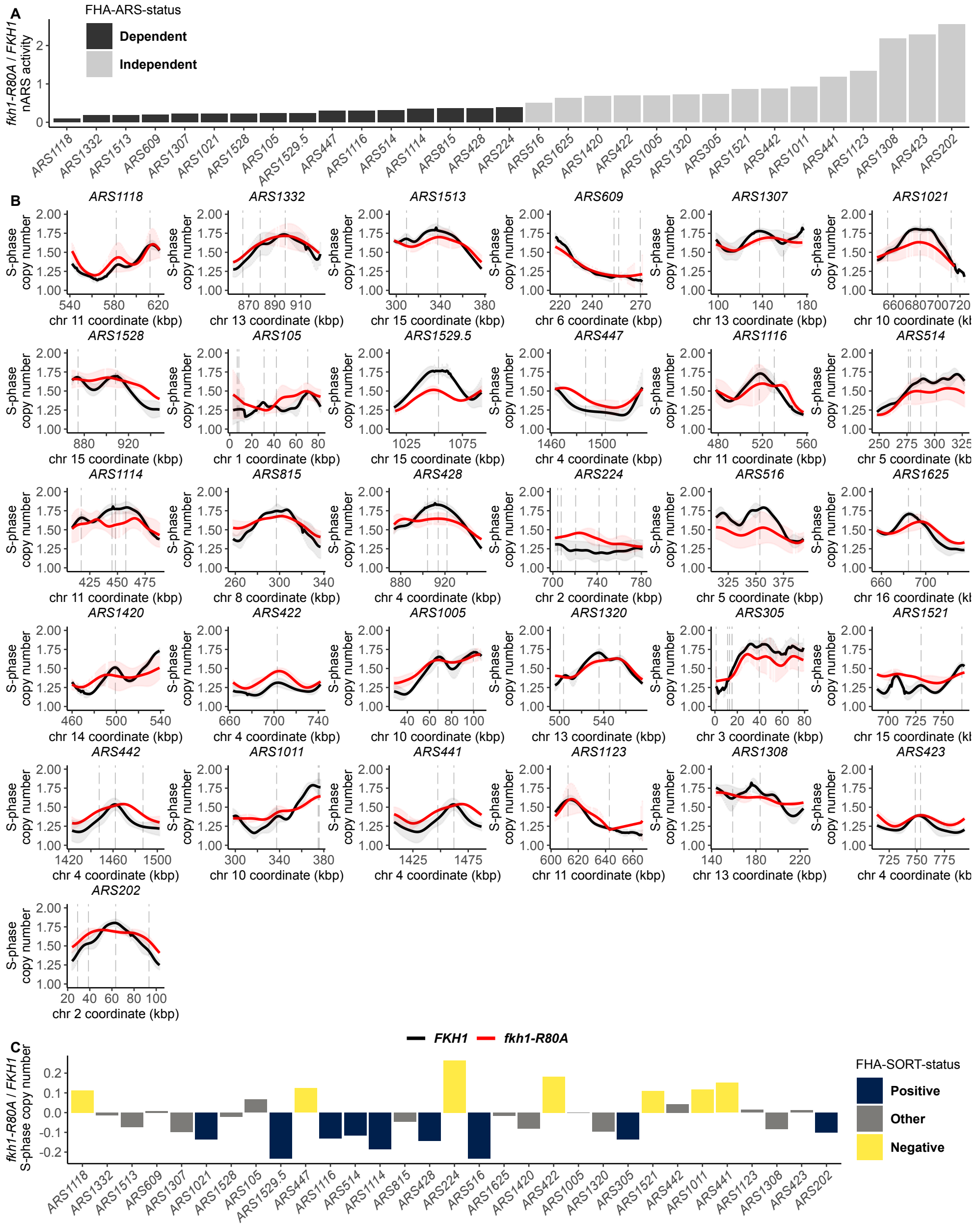


**Figure S3. Origin-by-origin assessment of the SortSeq data generated by the FHA-ARS-regulated origin groups.** **(A)** Ratio of ARS activity in *fkh1R80A* cells to *FKH1* cells (y-axis) for indicated origins (x-axis). FHA-ARS-dependent origins indicated by black bars while FHA-ARS-independent origins indicated by grey bars. **(B)** Close-up of S-phase copy number scans for FHA-ARS-regulated origins in *FKH1* (black) and *fkh-1R80A* (red) cells. **(C)** The *fkh-1R80A*/*FKH1* S-phase copy number ratio determined for each FHA-ARS-regulated origin color coded based on its FHA-SORT assigned status. FHA-ARS-regulated origins were defined initially based on ORC-origin binding mechanisms and subsequently parsed by ARS activity. Neither *ARS609* nor *ARS224*, each an FHA-ARS-dependent origin, qualified as FHA-SORT-positive. Note that neither origin is active even in FKH1 cells on the chromosome. Thus ORC-origin binding mechanisms might not play a major role in their chromosomal activity, yet might have a large impact on origin activity in a more isolated situation, such as on a plasmid.

#### Figure S4

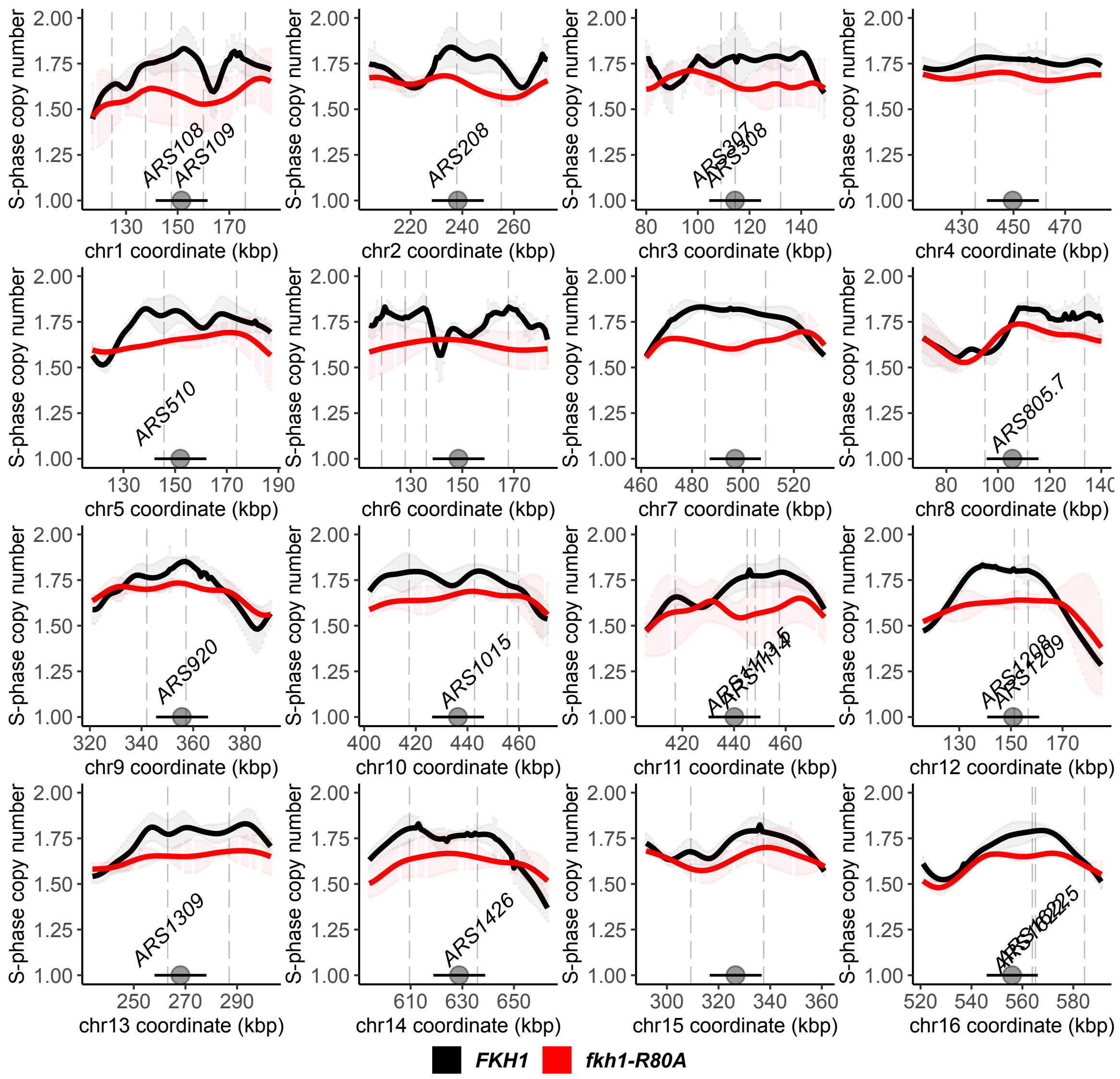


**Figure S4: Normalized S-phase copy number scans across yeast centromeres.** Based on the stringent definition of Cen-associated origins used here, *CEN4*, *CEN6*, *CEN7*, and *CEN15* do not contain CEN-associated origins per our definition. Copy numbers are presented as means and 95% confidence intervals from three *FKH1* (black) and two *fkh1-R80A* (red) replicates, as discussed in the main text.

### Figure S5

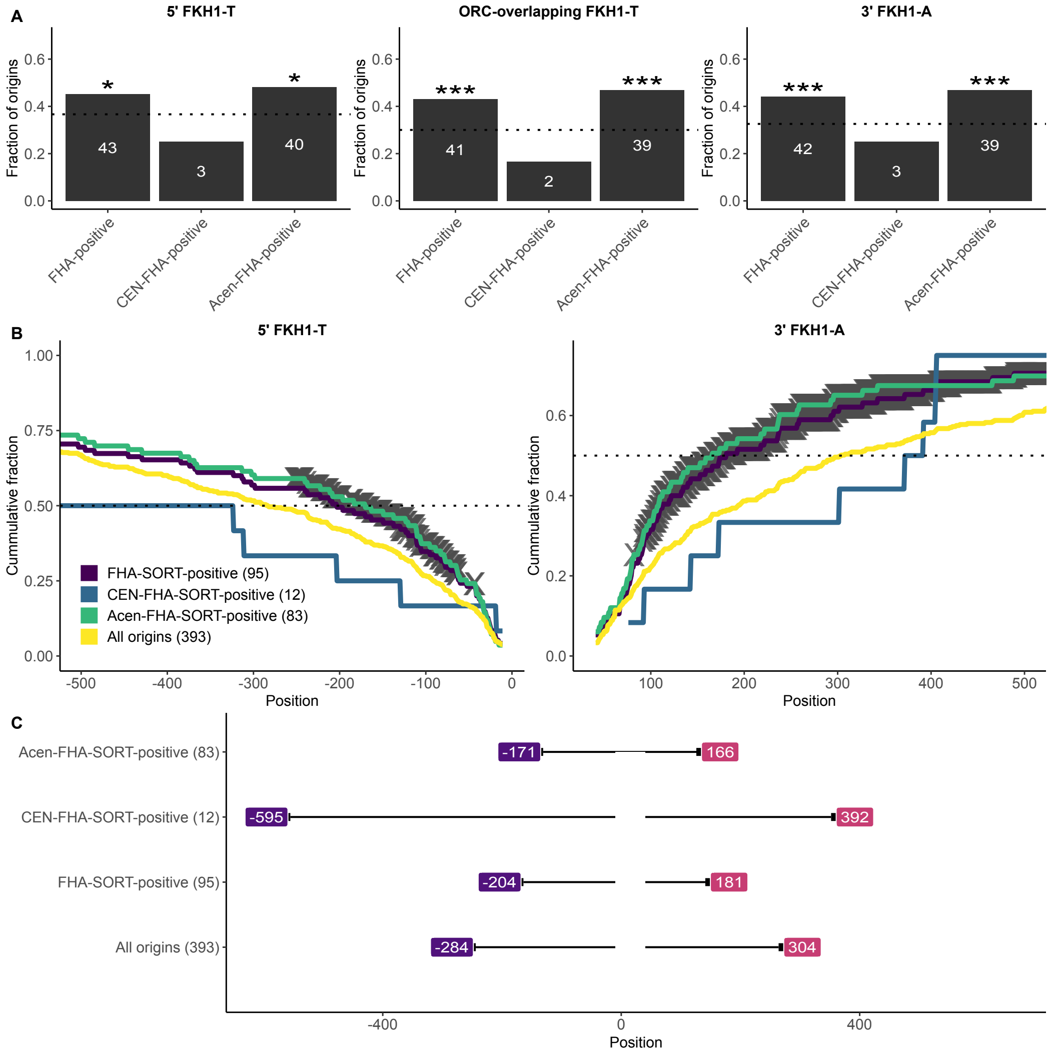


**Figure S5: Analyses of FKH1 motifs at FHA-SORT-positive Cen-associated origins.** The analyses in Figure 4, main text, were applied to Cen-associated origins that were FHA positive. **(A)** The fraction (y-axis) of origins within the indicated group (x-axis) containing at least one match to the indicated motif within the indicated origin regions for the relevant FHA-regulated origin groups classified by SortSeq. "5' FKH1-T" queried nucleotides -150 through -11 for FKH1-T motif matches, while "3' FKH1-A" queried nucleotides +41 through +150 for FKH1-A matches. The "ORC-overlapping" region encompassed nucleotides -10 through +40 and was queried for FKH1-T motifs. The horizontal line indicates the fraction of all confirmed origins (n=393) that contained a match to the queried FKH motif in the origin region under assessment. The enrichment or depletion of a given motif in any given origin group was challenged against the fraction of that motif in all confirmed origins using the hypergeometric distribution function. Significant P-values are denoted by asterisks (*, $P < 0.05$; **, $P < 0.001$; ***, $P < 0.0001$). In these analyses, 150 bp regions 5' and 3' of the ORC site were queried. **(B)** The cumulative fraction of origins (y-axis) in the indicated groups containing a 5'FKH-T (left) or a 3'FKH-A (right) after traversing the indicated number of nucleotides from the ORC site (x-axis). Nucleotide positions that reached P-value significance values of 0.01 are indicated by gray cross marks derived from hypergeometric distributions where at each position, the fraction of origins in the queried collection that contained a match by that nucleotide position was reached is compared to the fraction of all confirmed origins (n=393) that contained a match by the same position. **(C)** Summary of the 50% accumulation point for the analyses in (B).

### Figure S6

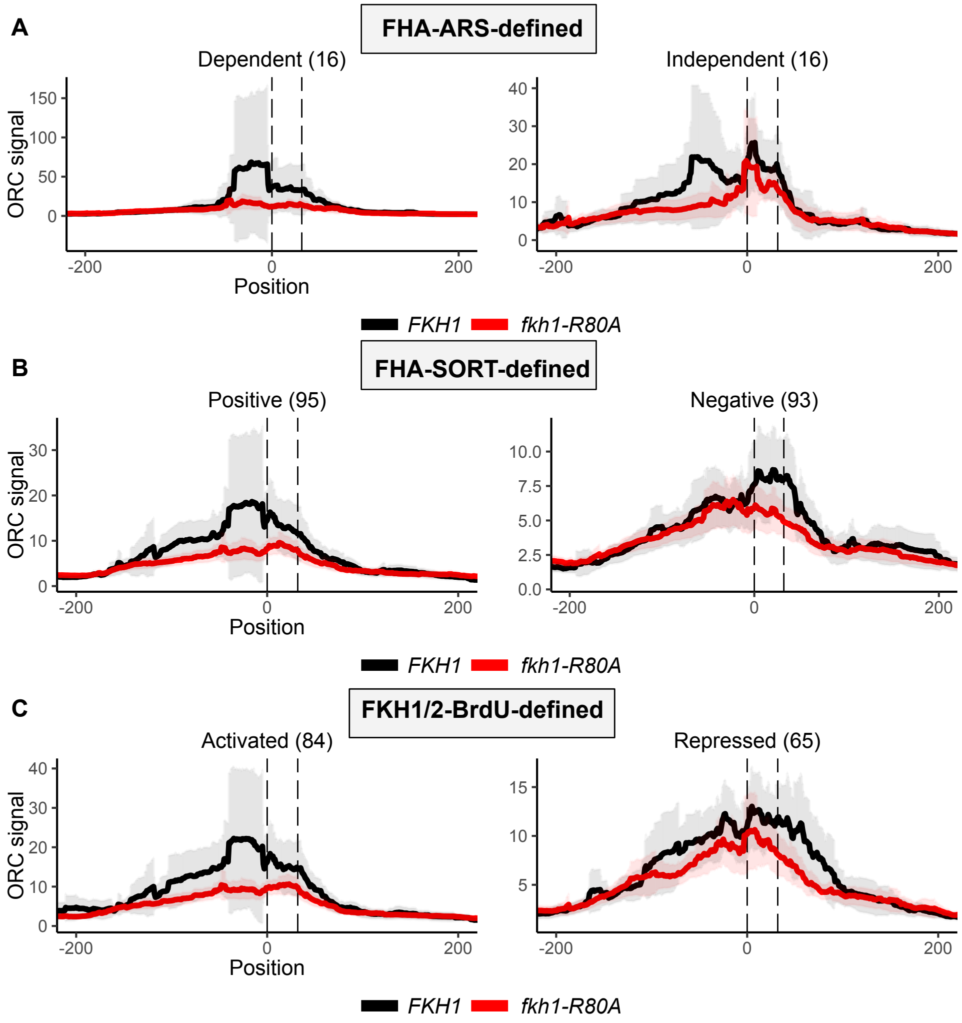


**Figure S6. Summarized internally scaled ORC binding levels at three types of origins.** Data are as presented in Figure 5, panels A, D, and G, but with the inclusion of 95% confidence intervals.

### Figure S7

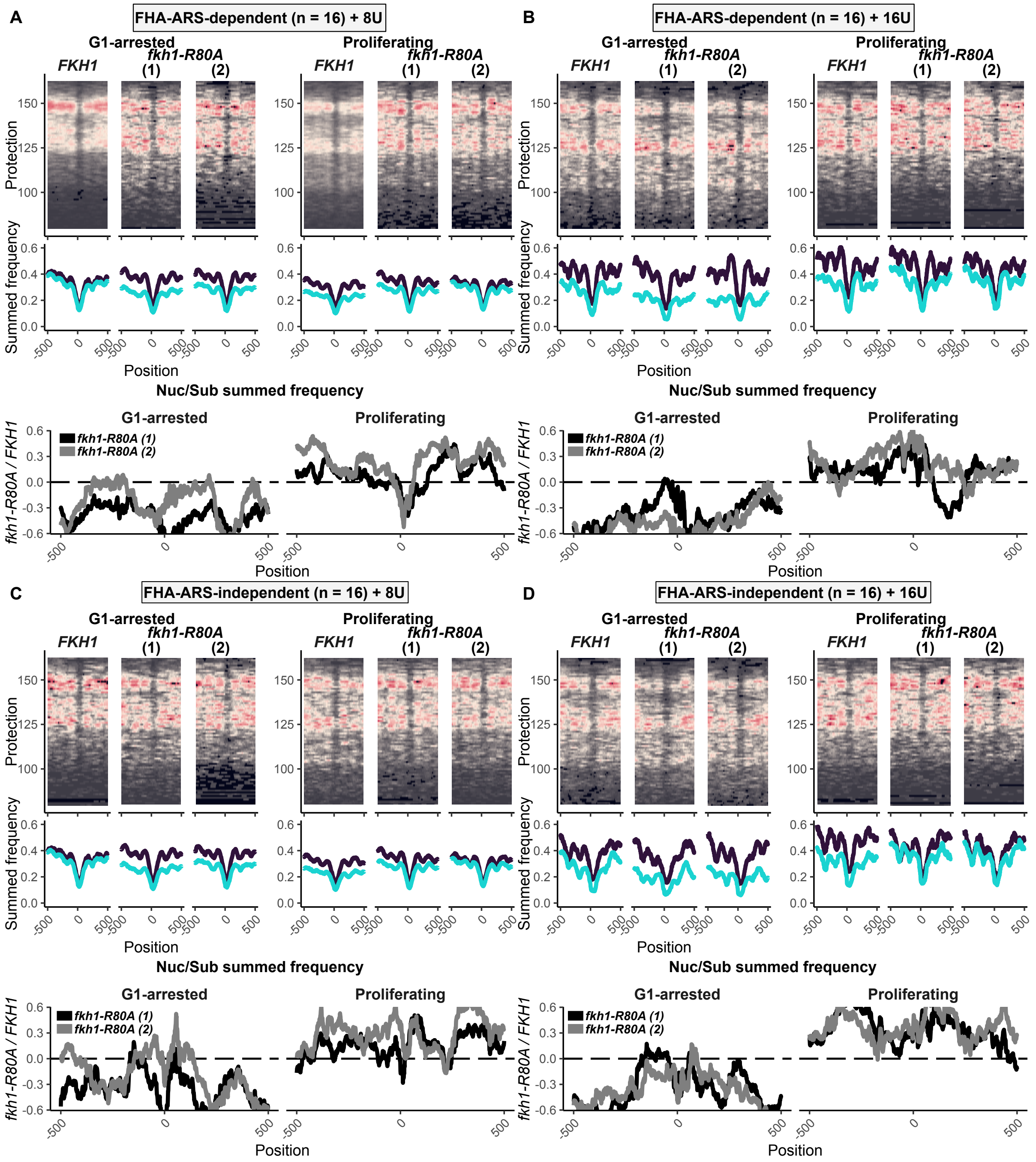


**Figure S7. The Fkh1-FHA domain promoted nucleosome stability similarly at FHA-ARS-dependent and FHA-ARS-independent origin groups.** The analyis of MNaseSeq data for all origins described in Figure 6 were applied to the contrasting groups of FHA-ARS-regulated origins as defined in Figure 2A.

### Figure S8

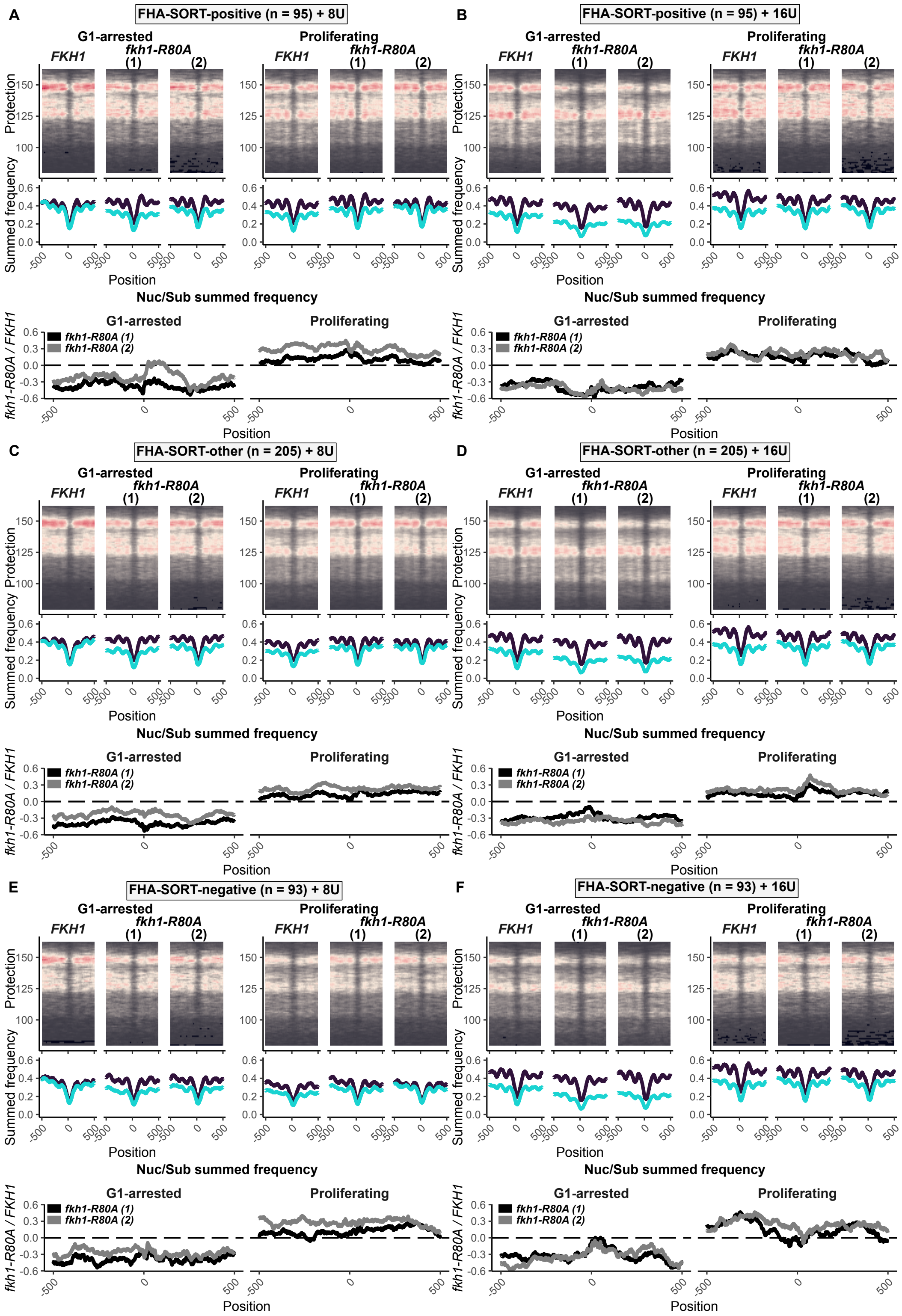


**Figure S8. The Fkh1-FHA domain promoted nucleosome stability similarly at FHA-SORT-positive and FHA-SORT-negative origin groups.** The analyses of MNaseSeq data for all origins described in Figure 6 were applied to the contrasting groups of FHA-SORT-regulated origins as defined in Figure 1 and Figure 2B.

### Figure S9

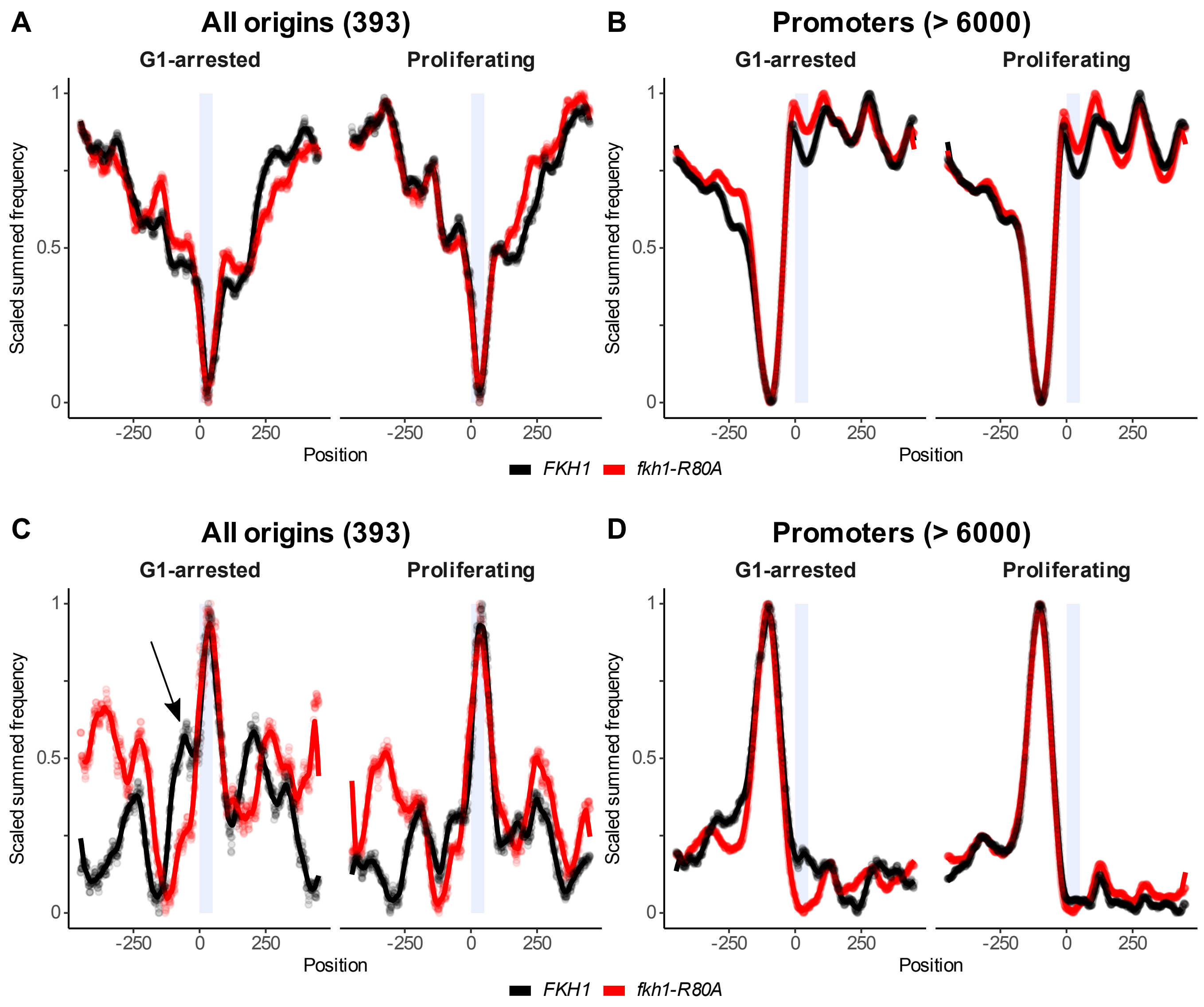


**Figure S9. Nucleotide-resolution, scaled nucleosome (A, B) or ORC (C, D) signals for all 393 origins or all 6000 promoters in *FKH1* and *fkh1-R80A* cells under G1-arrested or Proliferating conditions, as indicated.** The arrow in panel C indicates a G1-arrest specific and Fkh1-FHA-dependent 5' shoulder on the ORC peak.

#### Figure S10

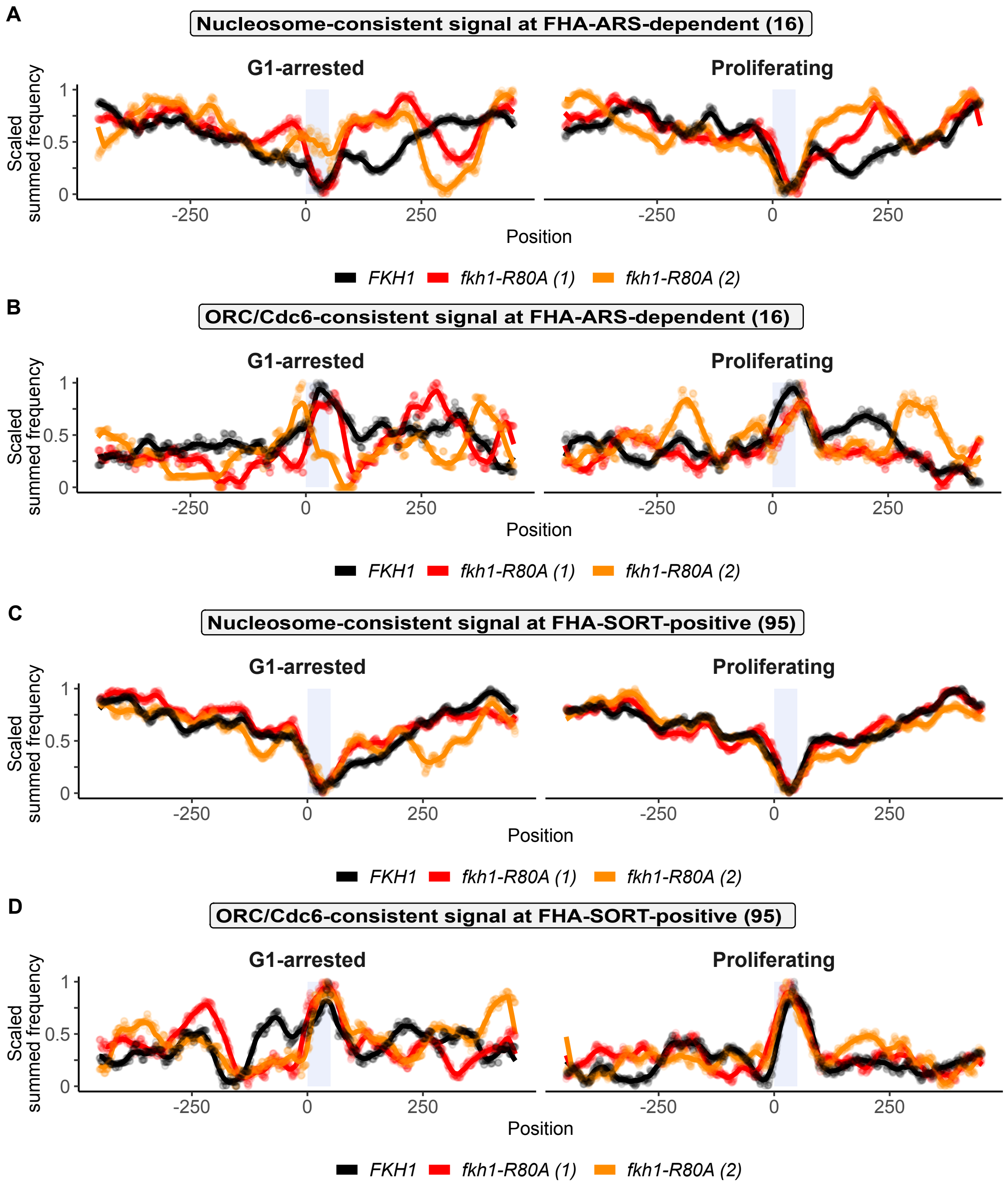


**Figure S10. The Fkh1-FHA domain promoted extended ORC signal at the FHA-SORT-positive origin group in G1-arrested cells as evidenced by two meiotically-derived *fkh1-R80A* replicates.** **(A)** Scaled nucleosome signals (generated from 8U MNase experiment) for the FHA-ARS-dependent origin cohort under G1-arrested and Proliferating conditions. Signal from *FKH1* and the two independent *fkh1-R80A* experiments are in black, red, and orange, respectively **(B)** As in (A) but with scaled ORC/Cdc6-consistent signals. **(C/D)** As in (A/B) but with FHA-SORT-positive origins.
